# DNA methylation is linked to the monoallelic expression of *MRP3*, a diatom mating type determining gene

**DOI:** 10.1101/2023.10.11.561864

**Authors:** Antonella Ruggiero, Svenja Mager, Francesco Manfellotto, Viviana Di Tuccio, Tomohiro Nishimura, Leslie Pan, Monia T. Russo, Remo Sanges, Maria I. Ferrante

## Abstract

Diatoms are unicellular microalgae widely distributed in aquatic ecosystems. In diatoms, sexual reproduction is needed to counteract cell miniaturization imposed by the rigid silica shell, and only small cells, below a species-specific size threshold, are competent for sex.

We performed a genome-wide Enzymatic Methyl-seq analysis in the heterothallic diatom *Pseudo-nitzschia multistriata* comparing cells of different size and opposite mating type (MT) to investigate potential epigenetic controls in life cycle transitions. We found an imprinting-like pattern of methylation at the sex locus: alleles of the gene responsible for the specification of the MT+, *MRP3*, are hypermethylated in MT- and differentially methylated in MT+, with transcription occurring only on the MT+ hypomethylated variant. The methylation pattern is overall stable over the *P. multistriata* life cycle. Absence of methylation in *MRP3* is necessary for its expression but not sufficient, since large non-sexual cells have the same methylation profile of small sexual cells but do not express *MRP3*, suggesting that additional controls are involved in the mechanism of sex determination.

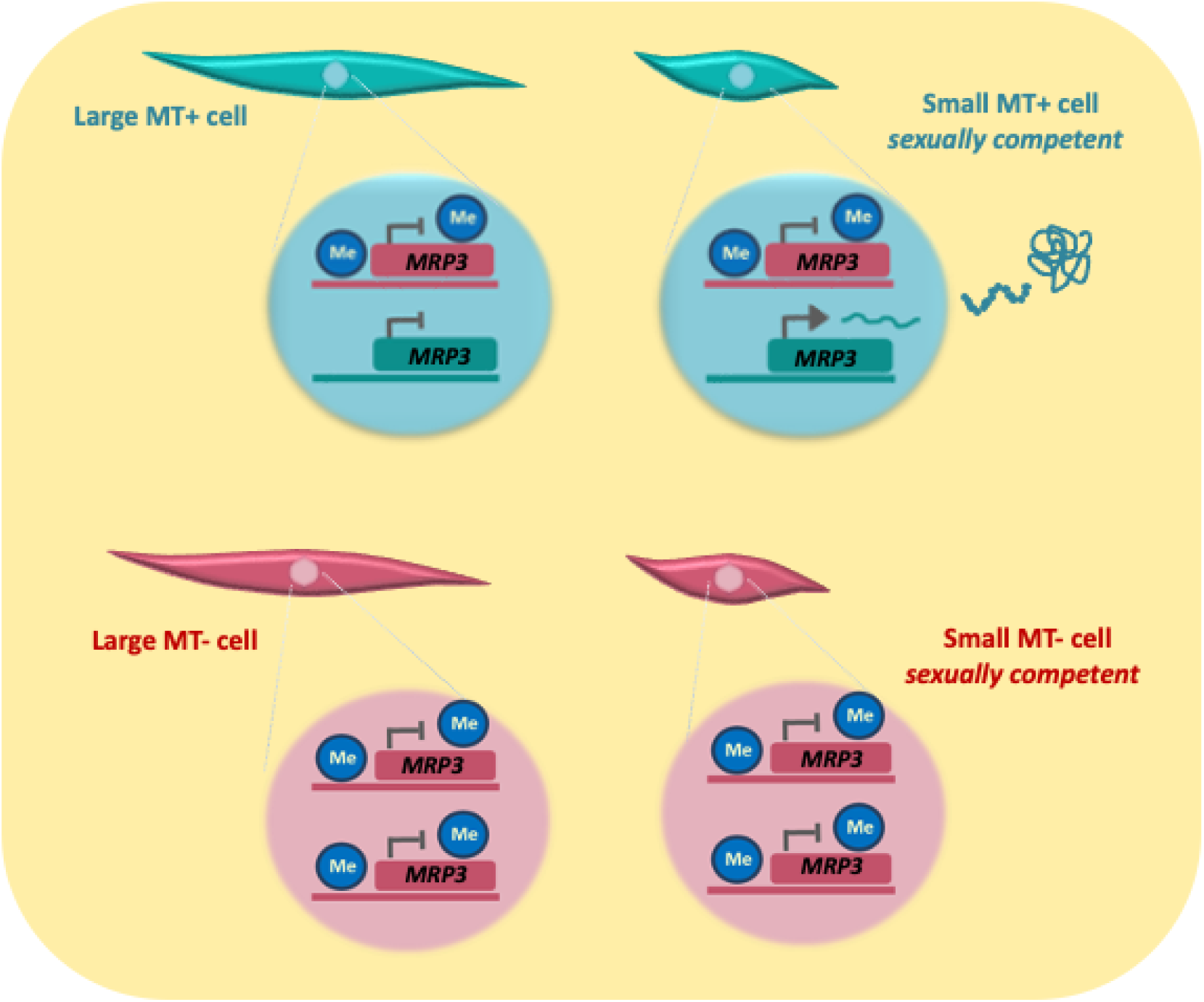

The methylation profile of the mating type determining gene *MRP3* of the diatom *Pseudo-nitzschia multistriata* is different between opposite mating types. Lack of methylation is necessary but not sufficient for *MRP3* expression in the sexually competent MT+ strains.

**Highlights:**

- - The mating type locus is differentially methylated in MT+ and MT-
- - DNA methylation is linked to MRP3 expression
- - DNA methylation does not play a role in the acquisition of sexual competence

## Introduction

Sex is widespread among eukaryotes, possibly evolving only once in the common ancestor (*Goodenough & Heitman 2014*), as suggested by the presence of meiotic recombination signatures in the genomes of all eukaryotic clades (*Speijer et al., 2015, Goodenough & Heitman, 2014*). In eukaryotic microbes the genes required for determining cell identity and directing sexual development are encoded in a specialized region of the genome known as the mating type (MT) determination region (*Lee et al., 2010, Heitman et al., 2015*), comparable to sex chromosomes. MT determination mechanisms evolved independently in different lineages and are extremely diversified. Unicellular organisms which display condition-dependent sex can shift between sexual and asexual reproduction alternating haploid and diploid phases in their life cycle. Asexual-sexual switch can be facultative in many groups, while it is mandatory in the case of most of the diatoms (*Chepurnov et al., 2004, Montresor et al., 2016*) which, among protists, represent one of the most diverse groups. Diatoms are photosynthetic microalgae generating about 20% of the oxygen produced on Earth, as much as the entire world’s rainforests (*Field et al., 1998, Saade & Bowler, 2009)*. Diatoms main feature is a bipartite, siliceous cell wall called frustule comprising two valves, the smaller hypotheca that must fit inside the larger epitheca *(Pickett-Heaps 1990*). The rigid frustule imposes a constraint during cell division, indeed at each mitotic division each of the two daughter cells inherits a maternal valve of different size with the consequence of a progressive cell size reduction of the population. This progressive miniaturization of the population can lead to extinction, and it is generally counteracted by the process of sexual reproduction (*Bilcke et al., 2022*). Therefore, in diatoms sexual reproduction has two functions, it introduces genetic diversity *via* recombination and it avoids population extinction. A unique property of diatom life cycles is the existence of a sexualization size threshold (SST): large, young cells are incapable of producing gametes, cells must be below a critical size to become competent for sexual reproduction when the right conditions are present (**Fig. 1**). Despite their ecological importance, very little is known on the molecular controls of the life cycle transitions and on the mechanisms involved in sex determination in diatoms (*Bilke et al., 2022*).

**Figure 1.**
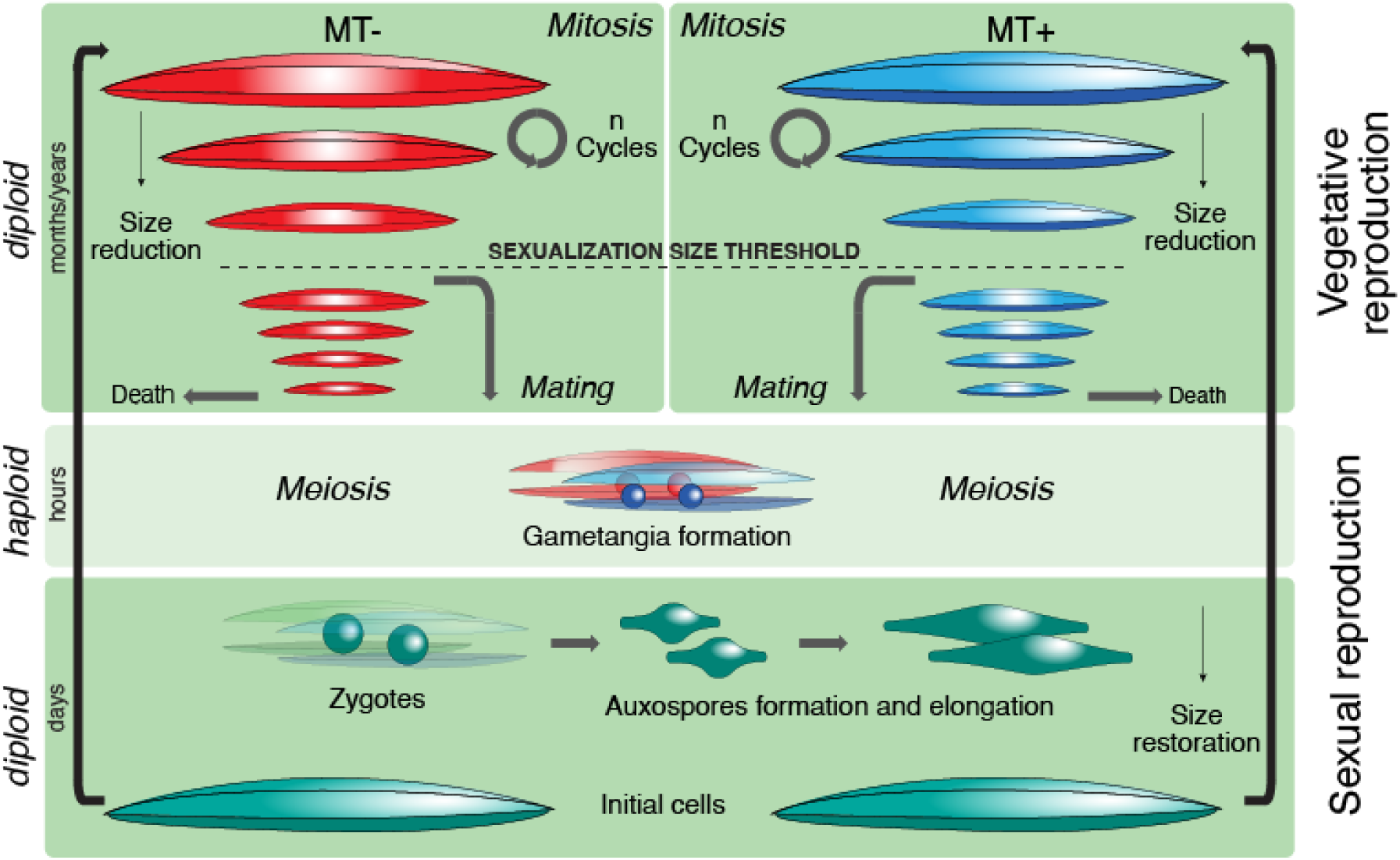
Schematic representation of the *P. multistriata* life cycle. Due to the presence of the rigid silica cell wall, mitotic divisions lead to the progressive reduction of the cell size. Below a species-specific size threshold (SST), cells can engage in sexual reproduction if they find a cell of opposite MT to pair with. Gametes are the only haploid stages in diatom life cycles. Zygotes are soft stages that can expand, forming an auxospore within which a large initial cell with a rigid cell wall is produced, thus restoring the maximum cell size and resuming the cycle.

Epigenetic mechanisms could in principle be involved in sexual development in diatoms because they provide the plasticity to combine genomic and environmental signals to influence gene expression through the regulation of chromatin structure and DNA accessibility (*Deans & Maggert 2015, Cheng et al., 2019, Feng et al., 2010, Piferrer 2013*). Cytosine methylation is a heritable DNA modification involved in the repression of transposable elements, gene expression, embryo development and sex determination in animals and plants (*Sun et al., 2022, Piferrer et al., 2021, Bräutigam et al., 2017, Edwards et al.2017).* In the process of DNA methylation a methyl group is transferred to the 5th carbon of cytosine. In mammals methylation is primarily found in the CpG context *(Goll and Bestor 2005)* while plants methylate in CG, CHG and CHH contexts, where H can be any nucleotide other than G (*Gruenbaum et al., 1981*).

DNA methylation has previously been investigated in diatoms (*Huff & Zilberman, 2014; Traller et al., 2016; Veluchamy et al., 2013, Gabed et al., 2022, Hoguin et al., 2023*), revealing the conservation of most of the enzymes required for the epigenetic machinery and confirming a major role of this process in the silencing of transposable elements (TEs).

Due to extensive background data, long-term observations and availability of molecular resources, including a sequenced genome *(Basu et al., 2017)*, the pennate diatom *Pseudo-nitzschia multistriata* is a suitable model to investigate the function of DNA methylation in sex onset and sex determination *(Ferrante et al., 2023)*. *P. multistriata* is a toxic species, able to produce the neurotoxin domoic acid *(Bates et al., 2018)*. It has a diploid genome and controllable life cycle, a heterothallic mating system with two mating types, MT+ and MT-, with a short generation time. Maximum cell size in *P. multistriata* is around 80 µm in length while sexual reproduction can occur only when cells reach the SST, which is around 55 µm *(D’Alelio et al., 2009)*. As in other pennate diatoms, sexual reproduction is triggered by the presence of a strain of the opposite MT. *P. multistriata* is the only diatom for which a master regulatory gene for sex determination has been identified, even though the molecular cascade is still under investigation. By comparing the expression profile of sexual competent cells of opposite MTs, MT specific genes were identified, <u>M</u>ating type <u>R</u>elated <u>P</u>lus *MRP1, MRP2* and *MRP3* expressed in MT+, and <u>M</u>ating type <u>R</u>elated <u>M</u>inus, *MRM1* and *MRM2,* expressed in MT-*(Russo et al., 2018)*. *MRP3* is the gene specifying the identity of the MT+, it is expressed in a monoallelic manner only in MT+ strains. Its overexpression in an MT-strain causes sex reversal: cells behave like MT+ and display an inverted trend of expression of the mating type related genes *(Russo et al., 2018)*. Different allelic variants of the region upstream of *MRP3* have been described (*Russo et al., 2018, Russo et al., 2021)*, characterized by polymorphic elements containing triplet repeats of different length or single nucleotide polymorphisms (SNPs) and a putative transposon, but their role in the regulation of the gene transcription remains to be clarified.

In this work we investigated the role of DNAmethylation in sexual reproduction on a genomic scale in a marine diatom, adding a piece of novel knowledge for the diversity and evolution of mating systems in unicellular organisms.

## Results and Discussion

To investigate the potential role of DNA methylation in life cycle transitions we profiled cytosine methylation of cells of opposite MTs above (large cells, L) and below (small cells, S) the SST **(Fig 1**) using Enzymatic Methyl-seq (EM-seq) analysis *(Vaisvila et al., 2021)*, across the entire genome and at single base resolution. In particular, we analyzed the whole genome of two sibling laboratory strains, MF9 (MT-) and MF11 (MT+), two independent wild strains from the Gulf of Naples (Italy), 1365-11 (MT-) and 1364-6 (MT+) and two independent wild strains from New Zealand, NZ5 (MT+) and NZ10 (MT-) **(Fig S1, *Table S1*).**

We first explored the *P. multistriata* methylation landscape, identifying methylation in the CG, CHG and CHH contexts. The vast majority of methylated cytosines were found within CpG dinucleotides, about ∼7-11% of CpGs were methylated in the 59 Mb *P. multistriata* nuclear genome, with slightly higher levels in the wild strains (9.3 and 9.8% for Gulf of Naples, and 10 and 11.5 % for New Zealand) compared to the lab strains (between 7.6 to 8.1%) (**Fig 2A, *Table S2***).

**Figure 2.**
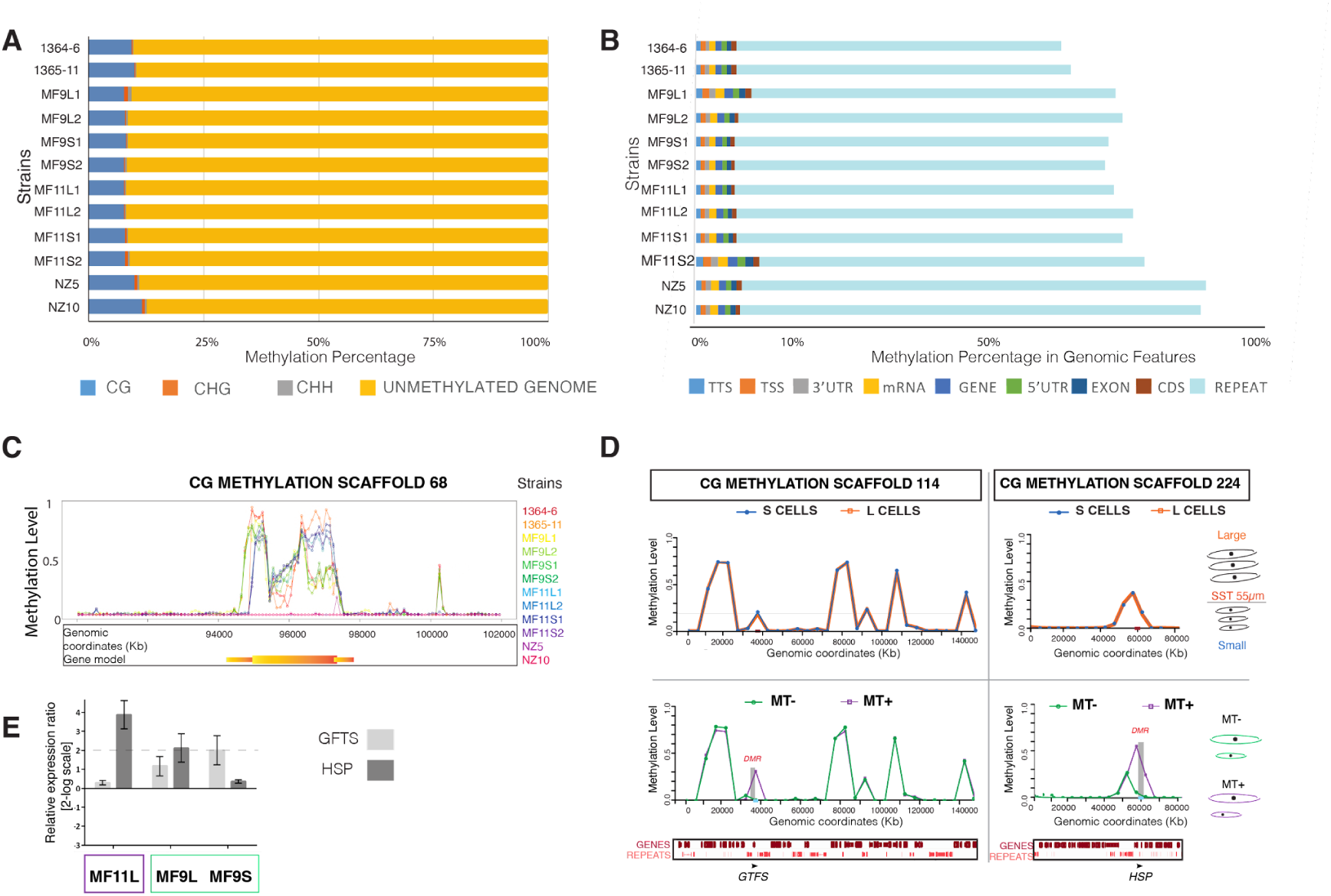
Methylation landscape in *P. multistriata* and expression levels of selected genes with a differential methylation profile. **A,** Percentage of methylation in the CG, CHG and CHH context in the different strains analyzed through EM-seq. **B**, Percentage of methylation in different types of genomic features: transcription termination site (TTS), transcription starting site (TSS), 3’ untranslated region (3’UTR), mRNA, gene, 5’untraslated region (5’UTR), exon, coding sequences (CDS), repeat. **C,** Methylation profile of different strains for gene PSNMU-V1.4_AUG-EV-PASAV3_0108790 on scaffold 68, the gene model is depicted below. **D**, Methylation profile on scaffolds 114 and 224 for large and small strains (top panels, orange and blue, respectively), and MT- and MT+ strains (bottom panels, green and purple, respectively). Gene models and repeats are indicated at the bottom The vertical gray bars mark the differentially methylated region (DMR). **E**, qPCR data for *GFTS* and *HSP*, MF11S is used as control condition (zero) to which the fold changes of the other samples refer. Error bars are calculated over three replicates.

In diatoms, the percentage of methylated cytosines varies markedly, with levels similar to what we found in *P. multistriata* (8.63% in *Fragilariopsis cylindrus, Huff & Zilberman, 2014*) but also much smaller (2.57% in *Thalassiosira pseudonana (Huff & Zilberman, 2014)* and 2.67% in *Haslea ostrearia (Gabed et al., 2022)*) or tremendously higher as for *Cyclotella cryptica* in which up to ∼60% methylation was found *(Traller et al., 2016)*. In agreement with what was found in other diatoms *(Huff & Zilberman, 2014 Traller et al., 2016; Veluchamy et al., 2013, Gabed et al., 2022*), most of the methylation was found in CG context while the global methylation level in CHG and CHH contexts was below 1% in all analyzed samples (**Fig 2A, *Table S2*).** About 70% of methylated CpGs were located within repeat regions, while we found only 0.8% in TSS and 0.7% in TTS (**Fig 2B)**, and between 1 and 1.5% in genes, where some genes were found to be highly methylated. The manually curated, filtered set of highly methylated genes (see Methods) contained 18 high-confidence genes **(*Table S3*)**, where methylation was scattered along the gene body (**Fig 2C)**.

To investigate possible changes in the DNA methylation profile during *P. multistriata* life cycle transitions, we looked for differentially methylated regions (DMR) between small and large cells, respectively competent and not competent for sex, and we also compared opposite MTs. No DMRs were found for CG or CHG context when comparing cells of different size (**Fig 2D**) and only 14 DMRs were found in the CHH context mainly located in repeat regions, except for two located in genes, an aldo-keto reductase and an uncharacterized protein ***(Table S4***).

Our results show that the global CpG methylation pattern is faithfully preserved during size reduction in the life span, suggesting that DNA methylation is unlikely to be involved in sexual competence acquisition. In contrast to this result, the comparison of opposite MTs revealed a small number of DMRs. In particular, we found 42 DMRs in the CHH context, with two of these DMRs being located in genes, an uncharacterized protein and an adenine methyltransferase (***Table S5***). In the CpG context we found 364 DMRs (***Table S6***). Among these DMRs, we found 17 high-confidence differentially methylated (DM) genes (***Table S6, S7***). There is not a univocal correlation between the methylation profile of a gene and its expression. For instance, in plants coding sequence methylation may have negative or positive effects on gene expression (*Kumar & Mohapatra 2021*). To explore the correlation between the methylation profile of the *P. multistriata* highly methylated genes and their expression levels, we randomly chose two genes, one annotated as a sterol 3-beta-glucosyltransferase (*GTFS*) and the other as a heat stress transcription factor A-1 (*HSP-A1*), with high methylation in MT+ strains MF11 and 1364-6 (**Fig 2D**), and evaluated their expression profile using RNA samples collected in parallel with the DNA samples used for EM-seq. Our results showed that the expression levels of *GTFS* and *HSP-A1* in MF11 were not substantially different when compared to MF9. Moreover, the level of *HSP-A1* was higher in MF11 large cells compared to MF11 small cells, even if the methylation profile for this gene was the same above and below SST (**Fig 2E**), indicating that there is not a consistent correlation between the methylation profile and the expression levels.

Interestingly, among the differentially methylated genes we found the sex-determining gene *MRP3 (**Tables S3*** **and *S7,* Fig 3A**), making the sexual locus epigenetically marked. Moreover, gene PSNMU-V1.4_AUG-EV-PASAV3_0020760, annotated as a methyltransferase (MTase), the only other gene in the gene-poor putative sex determination region, was also differentially methylated **(Fig 3B**). In fact, the entire 32 kb putative sex determination region had a distinct methylation profile with respect to flanking regions, most likely due to the presence of several repetitive sequences (**Fig S2**).

**Figure 3.**
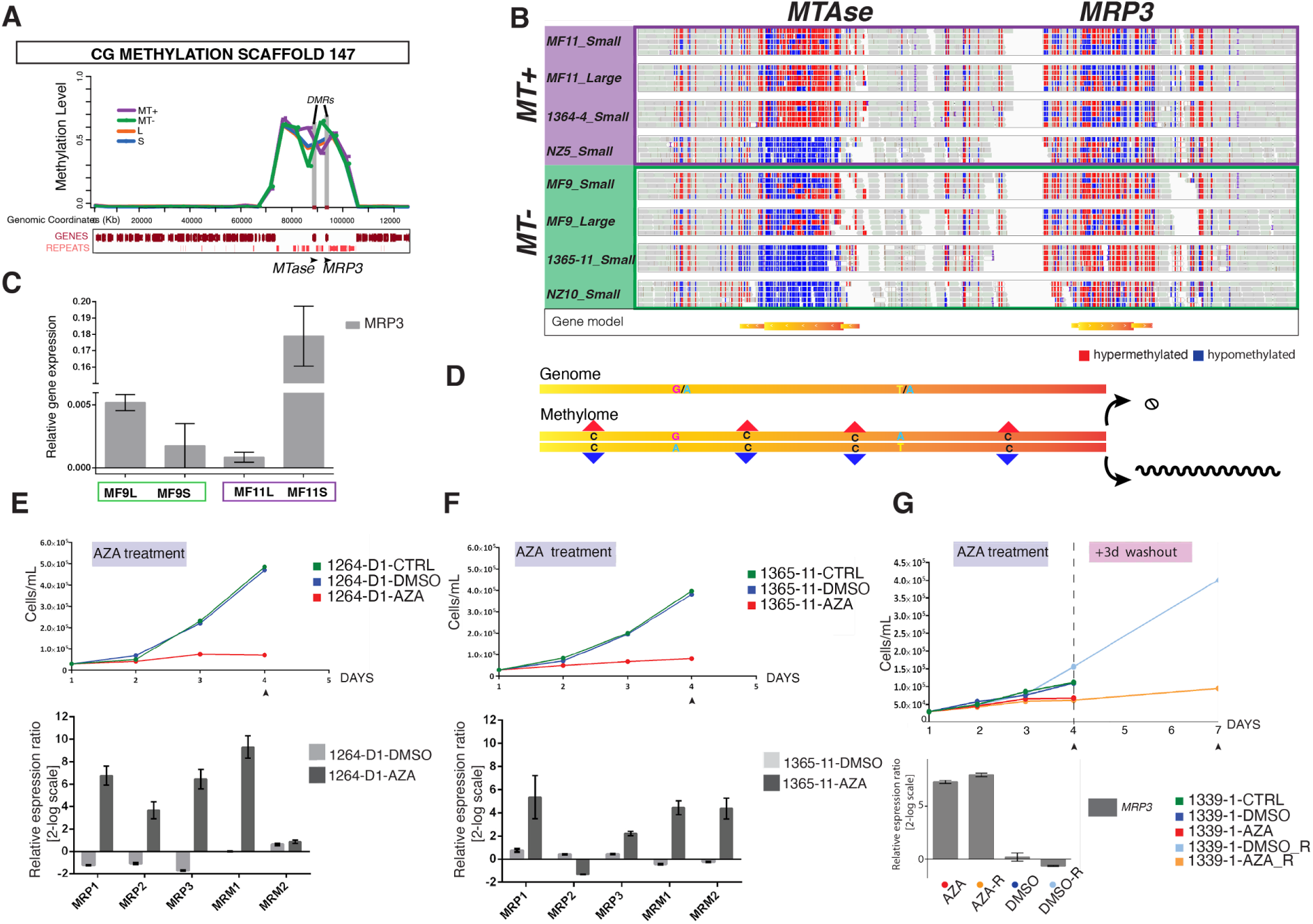
*MRP3* methylation profile and expression level following methylation inhibition. **A,** Methylation profile on scaffold 147 for MT+, MT-, large and small strains. Two DMRs are marked by vertical gray bars. **B,** IGV (Integrative Genome Viewer) EM-Seq genome visualization of a portion of scaffold 147 containing the *MRP3* and *Mtase* genes from all the strains used for this study (***Table S1***). Single reads can be visualized, blue squares represent hypomethylated regions, while red squares represent hypermethylated regions. The gene models are represented in yellow at the bottom. **C,** *MRP3* relative expression levels in large and small MF9 (MT-) and MF11 (MT+) cells. **D,** Schematic showing how SNPs allow to demonstrate that *MRP3* is transcribed from the hypomethylated allele (see **Fig. S3** for more details). **E-F,** Effect of AZA treatment on growth (top panels) and on the expression levels of MR genes (bottom panels) in two different MT-strains, 1264-D1 and 1365-11. The untreated sample is used as control condition and set as zero in the qPCR graphs. **G,** Growth curves and *MRP3* expression levels in the MT+ strain 1339-1 following AZA wash out, the untreated sample is used as control condition and set as zero.

We have shown that in *P. multistriata* the *MRP3* gene can have different alleles but only one, inherited exclusively by MT+ strains, is transcriptionally active *(Russo et al., 2018*). The second MT+ allele is not transcribed therefore the gene has a monoallelic expression. Structural genomic differences between MT+ and MT-strains in the promoter region of *MRP3* (*Russo et al., 2018, Russo et al., 2021)*, were thought to contribute to its expression regulation, although the present data suggest that the monoallelic expression may also be controlled by differential methylation.

We observed a different methylation pattern of the *MRP3* locus in MT+ and MT- (**Fig 3B**). In particular in MT-strains which do not express *MRP3* both alleles are hypermethylated (about 77% average CpG methylation) while in MT+ which express *MRP3* monoallelically only one allele is methylated (about 42% average CpG methylation) **(Fig 3B)**. By comparing SNPs on the reads, it is possible to deduce that the active *MRP3* allele is the one that is hypomethylated (**Fig 3D** and **Fig S3**). *MRP3* expression coincides with the acquisition of sexual competence, as shown by its restricted expression only in sexually competent MT+ cells **(Fig 3C)**, while the methylation profile is maintained between MT+ large and small cells **(Fig 3B)**. Our finding suggests that the difference in the methylation pattern between MT+ and MT-could play a role in sex determination rather than in sexual competence acquisition and that the *MRP3* expression is correlated to its methylation profile. We hypothesize that a transcription factor needed to activate *MRP3* expression becomes active at the SST switch in both MTs, but it can only access the unmethylated allele in the MT+, leading to restricted expression in the MT+ and subsequent activation of the MT+ genetic program. Additional experiments will be needed to test this hypothesis and identify putative candidates, such as RNA-seq and/or proteomics analyses comparing small versus large cells.

In MT+ strains isolated from the Gulf of Naples or from the Gulf of Mexico (*Russo et al., 2021),* the MT+ transcribed allele always bore some structural differences when compared to all other non-transcribed alleles (**Fig S4A** and **B)** *(Russo et al.,2018, 2021)*. The MT+ NZ5 strain deviates from this pattern as it has two structurally identical alleles (**Figs S4C**, **S5A** and **B**). We showed that in this MT+ NZ5 strain *MRP3* is upregulated with respect to an NZ MT-strain (**Fig S5C**), and it still retains a monoallelic expression (**Fig S5D**) and differential methylation on the two alleles (**Fig S5E**), strengthening the hypothesis that the different methylation is linked to the monoallelic expression. Indeed, nucleotide variations along the sequence allow to demonstrate that the expressed allele is the one with reduced methylation (**Fig S3D**). The MT+ specific allele of NZ5 appears overall identical in its structure to alleles typically found in MT-strains from the same and other geographical locations (compare **Fig S4B** with **S5A**). This implies that the distinctive repeats found in MT+ specific alleles from the Gulf of Naples and from the Gulf of Mexico (*Russo et al., 2018* and *2021*, **Fig S4B** bottom) are not responsible for the methylation status and/or the transcriptional control of the gene.

It is interesting to note that for the strains from the Gulf of Naples the MTase, the only other coding gene in the 32 kb MT determination region, also displays differential methylation (**Fig 3B**). In the MT-MF9 strain, the MTase appeared to be partially methylated, however this was not the case for the other two MT-strains analyzed (**Fig 3B**). In MF9, the MTase expression was found to be biallelic (**Fig S6A**), indicating that partial methylation for this gene does not correlate with selective expression from one of the two alleles. Moreover, the MTase expression does not appear to be linked to the MT nor to a specific combination of allelic variants (**Fig S6B and C**), suggesting that it is not involved in the sex determination cascade.

In eukaryotes four active DNA methyltransferases (DNMTs) families have been described *(Greenberg & Bourc’his 2019)*. As for other diatoms, *P. multistriata* lacks the DNMT-1 paralogue protein, required for methylation maintenance, probably supplied by DNMT5 which is involved in this process in a parasite fungus *(Huff & Zilberman 2014, Bewick et al 2019)*. *P. multistriata* also lacks *de novo* methyltransferase paralogue to DNMT3, as reported for *Pseudo-nitzschia multiseries (Hoguin et al., 2023)* (***Table S8***). We inhibited global methylation by treating MT-strains with the well-known DNA methyltransferase inhibitor, 5-azacytidine (5-AZA), which is incorporated into DNA as cytidine analogue that cannot be methylated *(Jones & Taylor, 1980*). If the lack of expression of *MRP3* in MT-cells is linked to the presence of methylation on both alleles, inhibiting the methylation should lead to increased *MRP3* transcription, which should in turn trigger upregulation of the MT+ specific genes and downregulation of the MT-specific genes, as observed in a transgenic MT-strain overexpressing exogenous *MRP3 (Russo et al., 2018)*.

The treatment with 5-AZA significantly reduced the growth rate of *P. multistriata* strains (**Fig 3E** and **F**, top panels). However, the overall cell morphology was not affected and cells were viable. We assessed the expression levels of MR genes in two MT-5-AZA-treated strains by qPCR (**Fig 3E** and **F**, bottom panels). Moreover, to discriminate against an unspecific effect due to the 5-AZA treatment, we used as negative controls genes that resulted not methylated and not linked to sexual reproduction, showing no changes in their expression (**Fig S7**). We found an overall upregulation of MT related genes due to global demethylation. *MRP3* and *MRP1* were consistently upregulated in treated MT-with respect to the untreated MT-, while *MRP2* was upregulated 6-fold in one experiment but it did not change in the other. Moreover, contrary to our previous findings in transgenic MT-cells overexpressing an exogenous MRP3 protein *(Russo et al., 2018)*, the expression of the MT-related genes *MRM1* and *MRM2* was not decreased but *MRM1* was upregulated in both cases while *MRM2* either did not change or was induced. It is important to note that *MRP1*, *MRP2* and *MRM2* do not appear to be methylated (**Fig S8A-B-D**), suggesting that their expression changed as a consequence of *MRP3* activation. On the other hand, we cannot conclude anything regarding the methylation status of *MRM1*, since there are no reads in the EM-seq data for the genomic region containing *MRM1* (**Fig S8C)**, perhaps due to the fact that this gene lies at the edge of the containing scaffold and must be in a region difficult to sequence. Whether 5-AZA directly affects *MRM1* expression therefore remains to be determined. 5-AZA strongly impacts cell physiology, neither MT+ nor MT-are able to mate following the treatment so we could not assess possible sex reversal.

We also followed *MRP3* expression after removing 5-AZA by washing cells and replacing the medium, finding that MT-cells resumed growth and could still express *MRP3* after seven days in drug-free medium (**Fig 3G**). These data suggest that the MT-*MRP3* methylation pattern is not restored during mitotic divisions.

Epigenetic controls in the sex determination regions have been reported for many species, from the most widely known human X chromosome inactivation to other diverse examples (*Muyle et al., 2021)*. Allele-specific hypermethylation of the SDR on the X chromosome was recently reported in channel catfish, where it silenced the *hydin-1* gene expression in genetic females *(Wang et al., 2022)*. While the hypothesis that methylation might play a direct role in sex chromosome evolution is controversial *(Pifferer, 2021)*, there are several examples reporting differential methylation on sex chromosomes and our findings represent the first report for a diatom, suggesting that *P. multistriata* harbors a primordial sex determining region, gene poor, repeat rich and with a distinctive methylation profile. *P. multistriata* is the only diatom in which a MT determination locus has been described so far, but current initiatives (https://jgi.doe.gov/csp-2021-100-diatom-genomes/) will provide genomic data on a larger set of diatoms, enabling further investigations on the evolution of sex determination systems and conservation of the genetic and epigenetic controls.

Interestingly, comparison of data from the lab strains above and below the SST and within replicates indicated that the general methylation profile is faithfully conserved when cells are kept under the same environmental conditions. It will be interesting to explore variability of these profiles under changing external conditions.

As for the mechanisms that allow cells to perceive their size and activate competence for sex, it will be interesting to explore transcriptional controls as well as other layers of epigenetic regulation, such as histone marks, and the general chromatin accessibility through dedicated approaches, such as ATAC-seq.

## Conclusions

Our study revealed a differential methylation pattern for the *P. multistriata* sex determining master gene between the two MTs, indicating that its expression is to some extent regulated by DNA methylation. In this specific case *MRP3* gene body methylation is linked to repressed transcription, representing a good example of clear correlation in a very diverse and controversial field (*Dixon & Matz 2022*). We assigned for the first time a function to DNA methylation in a specific phase of the diatom life cycle. The methylation profile in MT+ does not seem to be linked to specific structural features of the *MRP3* genomic region, as evident from the differential methylation profiles of the two structurally identical alleles in a MT+ New Zealand strain. Moreover, similar profiles for the *MRP3* region between large and small MT+ suggest a role for methylation in regulating accessibility for an as yet unidentified factor, only expressed in small cells, to one of the two MT+ alleles. On the other hand, the global methylation pattern was overall preserved during the whole vegetative life of cells, with differences found mostly in the CHH context in intergenic regions, suggesting that DNA methylation is not a major regulatory mechanism in the switch to sexual competence at the SST. The whole genome analysis also showed that repetitive regions (transposable elements) are major targets of DNA methylation in *P. multistriata*, confirming previous observations in diatoms *(Tirichine et al., 2013, Trelleret al., 2016, Hoguing et al., 2023)*. Finally, the analysis of strains from a different geographical location allowed us to expand our knowledge of the existing variants for the MT determining region. Future improvements in the genome contiguity and refinement through the use of long reads sequencing technologies, coupled with novel datasets for additional strains, will allow us to address pending questions on different topics touched in this study, including the specific role of CHH methylation, the role of the genes affected by differential methylation, and the conservation of the MT determining region.

## Materials and Methods

### Strains, culture condition and experimental crosses

*Pseudo-nitzschia multistriata* strains were isolated from the Long Term monitoring station MareChiara (LTER-MC) in the Gulf of Naples *(D’Alelio et al., 2010)*, and from the coasts of New Zealand *(Nishimura et al., 2021)*. F1 strains were manually isolated from experimental crosses of strains from LTER-MC (***Table S1***). Individual *P. multistriata* cells were isolated from the water sample with a micropipette and grown in f/2 medium supplemented with silicates *(Guillard, 1975)* at 18 °C, 50 μmol photons m^−2^ s^−1^ and a 12 L: 12 D h photoperiod. To test sex compatibility and assess the strains MT, they were crossed with reference MT- and MT+ strains isolated at LTER-MC using the same procedure described in *(Scalco et al. 2016*). Crosses were carried out in 6-well culture plates (Corning® Costar®) filled with 3 mL of f/2 medium. Exponentially growing cultures of pairs of strains were inoculated at a final cell concentration of ∼ 6×10^3^ cells ml^-1^. Culture plates were incubated at the culture conditions indicated above for five days and inspected daily to check for the presence of sexual stages. Single large F1 cells were isolated with a micropipette.

The isolated F1 initial cells (around 75 µm in length) were grown under the mentioned culture conditions for six months to reduce their size. F1 large (L, above 55 µm) and small (S, below 55 µm) cells were isolated from the mixed-size culture using a micropipette. Cultures of large and small F1 cells for EM-seq were collected in duplicate and processed for DNA and RNA extraction.

### DNA extraction

Before extraction of DNA, strains were grown in f/2 medium supplemented with antibiotics, as described in (doi.org/10.17504/protocols.io.bgudjws6). The absence of bacteria was tested as described in (dx.doi.org/10.17504/protocols.io.btt5nnq6).

Genomic DNA was extracted following the protocol described in (dx.doi.org/10.17504/protocols.io.byk7puzn) and quantified using Qubit® 2.0 Fluorometer (Life Technologies) while integrity was assessed by 1% agarose gel electrophoresis.

### Mating type determining region characterization

PCRs for mating type prediction were performed using WonderTaq (Euroclone SPA) and PCRs to define the structure of the *MRP3* alleles were performed using Q5® High-Fidelity DNA Polymerase (New England Biolabs, Massachusetts, USA), according to the manufacturer’s instructions using the primers reported in *Russo et al. (2018 and 2021*). The *MRP3* (GenBank ID MW116958) amplicons were obtained with a couple of primers that amplified the complete coding sequence, FMrp3Eco and RMrp3Sma (*Russo et al., 2018*). Sanger sequencing was performed on an Applied Biosystems (Life Technologies, Thermo Fisher Scientific, Waltham, MA, USA) 3730 Analyzer 48 capillaries using the primers reported in *Russo et al. (2018 and 2021*).

### Enzymatic methyl sequencing library preparation and sequencing platform

Libraries for 12 samples (***Table S1***) were prepared using the NEBNext Enzymatic Methyl-seq Kit (New England Biolabs, Massachusetts, USA) following the manufacturer’s large insert libraries protocol. Based on the quality assessment, the samples were standardized depending on the available amount of DNA in a volume of 50 µl, including a spike-in of pUC and Lambda DNA provided with the kit, as a control for methylation efficiency as per manufacturer’s protocol. The samples were fragmented using the Covaris S2 system, with settings to achieve an average fragment size of 350-400 bp. The samples were individually barcoded during the PCR using the PCR cycle numbers as per the manufacturer’s protocol depending on used input. The libraries were quantified using the Qubit HS DNA assay following the manufacturer’s protocol. The quality and molarity of the libraries was assessed using Agilent Bioanalyzer with the DNA HS Assay kit as per the manufacturer’s protocol. The assessed molarity was used to equimolarly combine individual libraries into pools for sequencing. All samples were sequenced with Illumina NextSeq 550/2000 platform (Illumina, San Diego, CA, USA) using a 300 cycle kit and a read-length of 150 bp paired-end reads.

### Sequencing data processing and methylation extraction

The raw reads were quality checked with fastqc v0.11.9 *(Andrews, 2010)*. Mapping and methylation extraction and quantification was done with Bismark v0.23.0, using bismark_genome_preparation with default parameters to index the reference genome *(Basu et al., 2017)* and then reads were aligned with default parameters and the resulting bam files deduplicated. Validation of the files was done with the ValidateSamFile program v2.26.6 of Picard (Broad Institute, 2019) and mapping statistics were called with qualimap v2.2.2 *(García-Alcalde et al., 2012)*. Methylation was extracted with bismark_methylation_extractor with--comprehensive option.

To determine the average methylation level genome-wide as well as in different genomic features (CDS, exon, 5’UTR, genes, mRNAs, repeats, 3’UTR, TSS and TTS), the methylation level of each cytosine of a certain context within a feature was determined (methylated reads divided by total reads, only considering sites with at least 5 reads) and then the average calculated. Raw data are available in ArrayExpress with accession number PRJNA889340 (data will be published upon paper publication)

### Differentially methylated regions determination

Differentially methylated regions (DMRs) were determined with the R package DMRcaller v1.26.0 and Bismark cytosine reports. DMRs were determined for comparison of small cells versus large cells as well as MT-samples versus MT+ samples from the Gulf of Naples, Italy. Valid DMRs in CG context needed to have a p-value of ≤ 0.01, a minimum size of 50 bp, a cytosine count of 5, a minimum of 5 reads per cytosine and a minimum proportional difference in methylation level of 30% between the comparing groups (small vs large cells, MT- vs MT+ samples). As suggested for CG DMRs by *Catoni et al., 2018* noise filter method with triangular kernel for noise filtering was used and the Score test as statistical test. For DMRs in CHG context the same parameters were used with the only modification of allowing DMRs with a difference minimum of 20% between the compared conditions. Finally, for CHH context DMRs, the bins method was used (with a bin size of 100) and minimal methylation difference was reduced to 10% difference between compared conditions.

For the comparison of large versus small cells, an additional analysis was done for DMRs in CG context, allowing DMRs with only a single differentially methylated cytosine between the two conditions and a minimal proportion difference required of only 10%.

### Determination of high confidence differentially methylated and highly methylated genes

We wanted to investigate genes that overlapped with a DMR as well as genes with high methylation level in CG context. CG methylation levels in each gene and for each sample were determined via ViewBS v0.1.11 MethHeatmap command. Since global CG methylation levels in the samples ranged between 7.6 and 9.8%, we considered only genes with a methylation level > 10% in all of the samples as highly methylated.

With bedtools v2.30.0 intersect command (applying –wa and –wb options), we determined DMRs overlapping with genes.

Since repeat regions were found to be globally highly methylated, we considered as highly methylated or differentially methylated only genes that were not overlapping with a repeat, assessed by visualizing each gene in the Integrative Genomics Viewer (IGV).

Moreover, since the *Pseudo-nitzschia multistriata* gene models are not always accurate, we also manually filtered out genes with a substantial discrepancy between the gene model structure and the corresponding RNA-seq reads available in the genome browser, in other words we discarded gene models that were not supported by the transcriptomics data available for the species (see RNA-seq tracks at http://bioinfo.szn.it/pmultistriata/).

### DNA Methylation players

DNMRT players in *P. multistriata* were determined by searching in the PLAZA Diatoms 1.0 platform (*Osuna-Cruz et al., 2020*) the gene IDs based on Table 1 in *Zhao et al., 2022*, which summarizes putative methylation players (proteins to set, maintain or remove cytosine methylation) in three different diatom species.

### DNA Methylation Inhibition

5-azacytidine (5-AZA) was prepared in 50% DMSO and 50% ice cold MilliQ water, and used at 20 μM final concentration. Lower doses were tested but no effect was observed neither on cell growth nor on gene expression levels. Stock solutions of 5-AZA were stored at -80°C; aliquots were thawed on ice prior to treatment to prevent drug instability and break down. *P. multistriata* cells were synchronized with prolonged darkness for 18 h. Following re-illumination, 5-AZA was added once a day for 3 days in the light cycle. Cell count was performed every day using the Malassez counting chamber. At the end of the treatment cells were harvested for gene expression analysis.

### RNA extraction, RNA sequencing and gene expression analysis

RNA extractions were performed as described in dx.doi.org/10.17504/protocols.io.261gen627g47/v1. RNA quantity was determined using a Qubit 2.0 Fluorometer (Life Technologies, Thermo Fisher Scientific, Waltham, MA, USA) and integrity using a 2100 Bioanalyzer System (Agilent Technologies, Santa Clara, CA, USA). RNA-seq library preparation and sequencing libraries for two samples (***Table S1***) were performed at the Genecore Facility of the European Molecular Biology Laboratory (EMBL) in Germany as described in *Annunziata et al. 2022*. Raw data are available in ArrayExpress with accession number E-MTAB-12676.

From 0.5 μg to 200 ng of total RNA was reverse-transcribed using the QuantiTect® Reverse Transcription Kit (Qiagen, Hilden, Germany). Gene expression levels were evaluated by qPCR using primers reported in ***Table S9***, *Russo et al., 2018* and *Russo et al., 2021*. The qPCR experiments were performed in triplicate in a ViiA 7 Real-Time PCR System (Applied Biosystems, Foster City, CA, USA) using Fast SYBR Green Master Mix (Applied Biosystems, Foster City, CA, USA), following manufacturer instructions. The reference genes used were *TUB-A* and *CDK-A* (Adelfi *et al*., 2014). The relative gene expression quantification was calculated with the following formula: relative expression ratio of the gene of interest is 2^−ΔCT^ with ΔCT = Ct_(target gene)_ minus CT_(reference gene)_.

Gene fold change relative expression was made following the Δ-Δ-Ct method. The results are the mean SD of at least three separate experiments, measuring each parameter by triplicate (n = 3).

## Acknowledgments

The authors wish to thank Enrico D’Aniello, Stefania Castagnetti and Tobias Gerber for the useful comments on the manuscript. We thank the Stazione Zoologica Sequencing and Molecular Analysis Center (CSAM) for Sanger sequencing and RNA quality assessments and the EMBL GeneCore service for EM-seq and RNA-seq. This work was supported by the Gordon and Betty Moore Foundation through Grant #7978. The New Zealand Ministry for Business, Innovation and Employment (Safe NZ Seafood contract No. CAWX1317) contributed some funding towards this study.

## Supplementary materials

**Figure S1.**
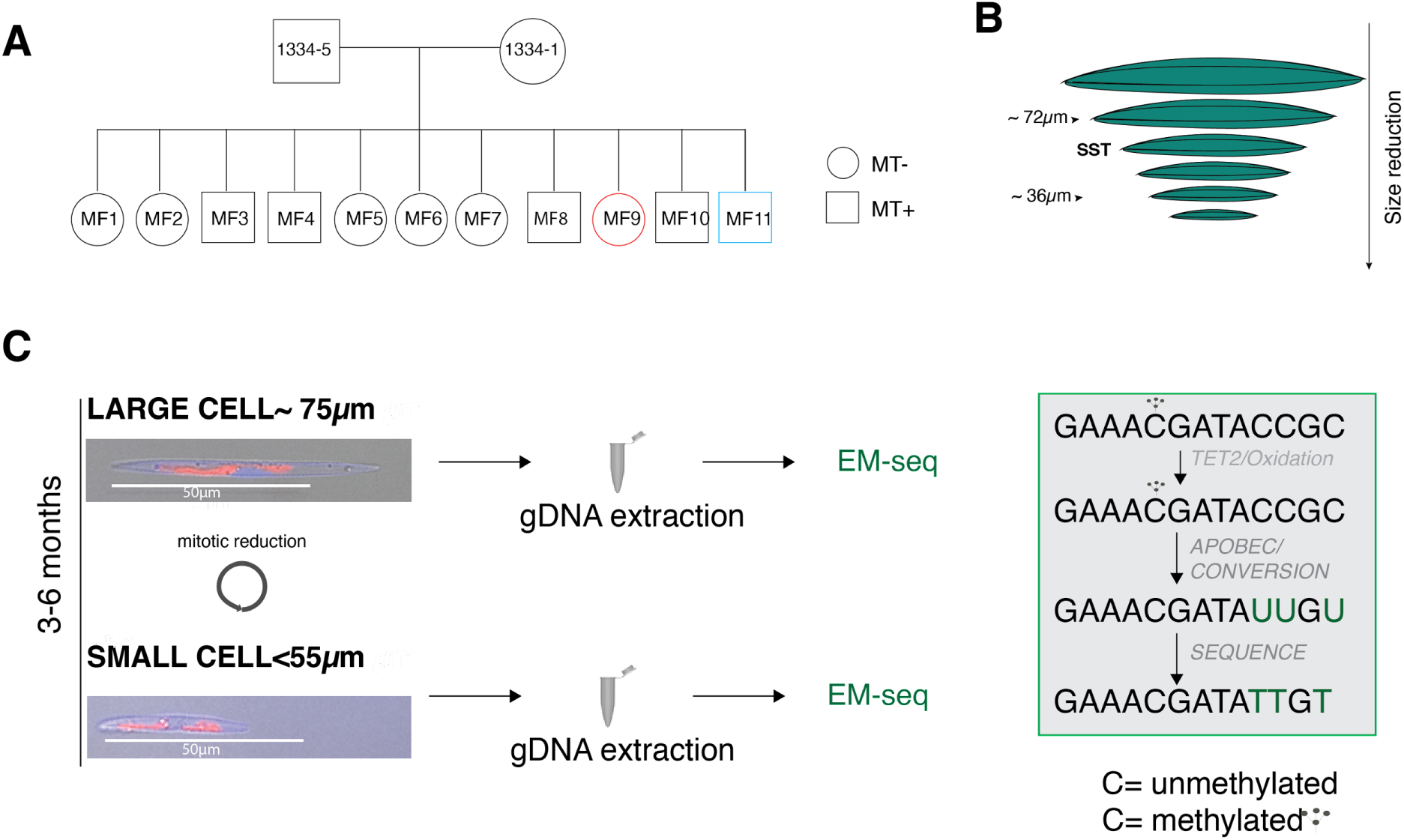
Experimental set up for genome wide methylation analysis of large and small *P. multistriata* cells. **A**, *P. multistriata* pedigree, showing one generation which includes two sibling strains of opposites MTs, MF9 and MF11, selected for the methylation analysis. Single large F1 cells were manually isolated and the established monoclonal cultures were grown for a few months to obtain small cells. **B**, Schematic showing cell size reduction with arrows indicating the approximate size of cells collected above and below the SST from each monoclonal culture. **C**, Large and small cells were grown under standard laboratory conditions and harvested at exponential growth phase. High quality gDNA was extracted and processed for the EM-seq analysis. On the right panel a representation of the EM-seq method is shown: enzymatic modifications convert Cs into Us, except when they are protected by methylation. In the micrographs, merged bright field and fluorescence visualization is shown, red is the chlorophyll autofluorescence in the chloroplasts, scale bar 50 µm.

**Figure S2.**
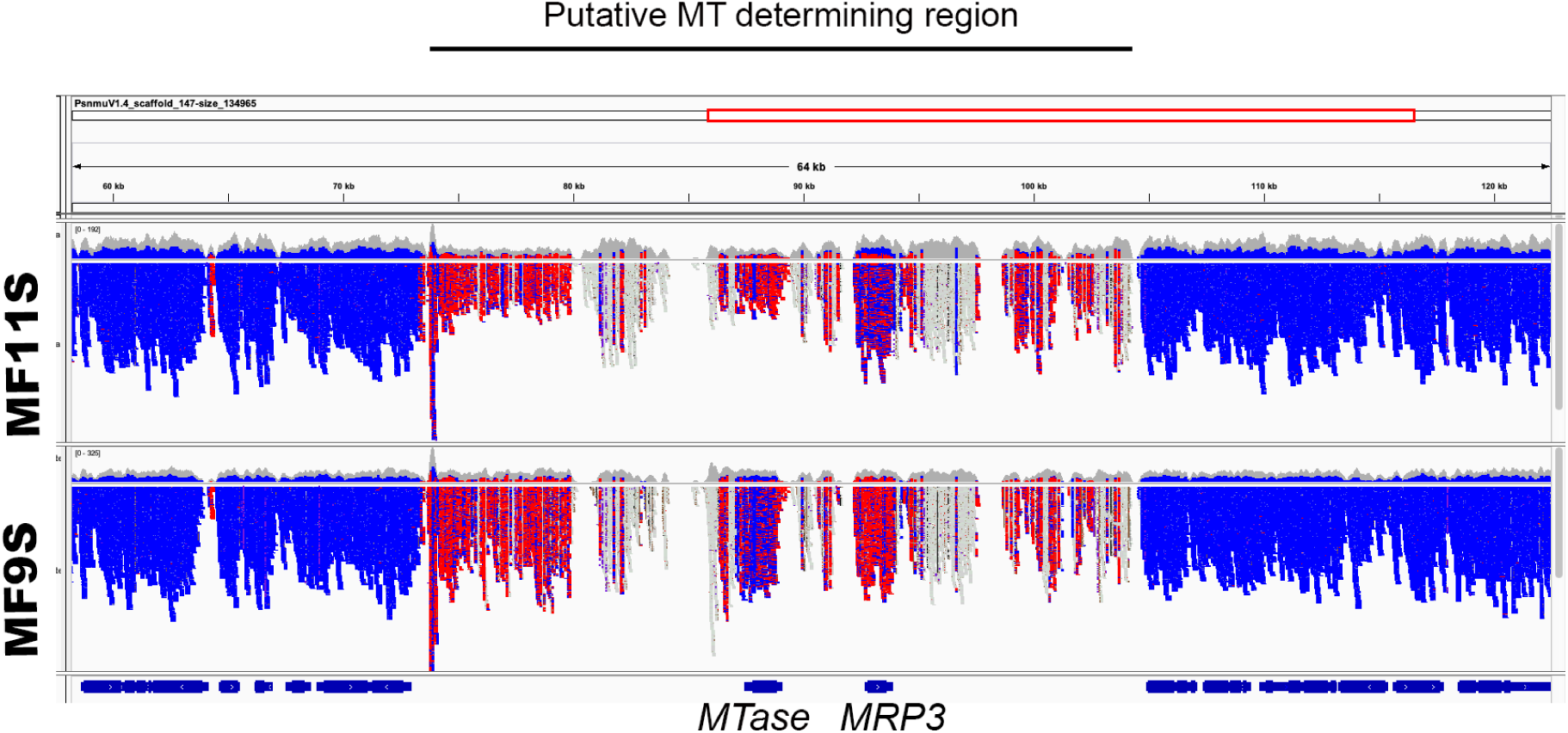
Methylation profile on scaffold 147. IGV visualization of the region between positions 58,188-122,460 for strains MF11S (MT+) and MF9S (MT-). The putative 32 kb mating type determining region displays a different methylation profile compared to the 5’ and 3’ flanking regions, which are mostly hypomethylated. Gene models are shown at the bottom as blue bars.

**Figure S3.**
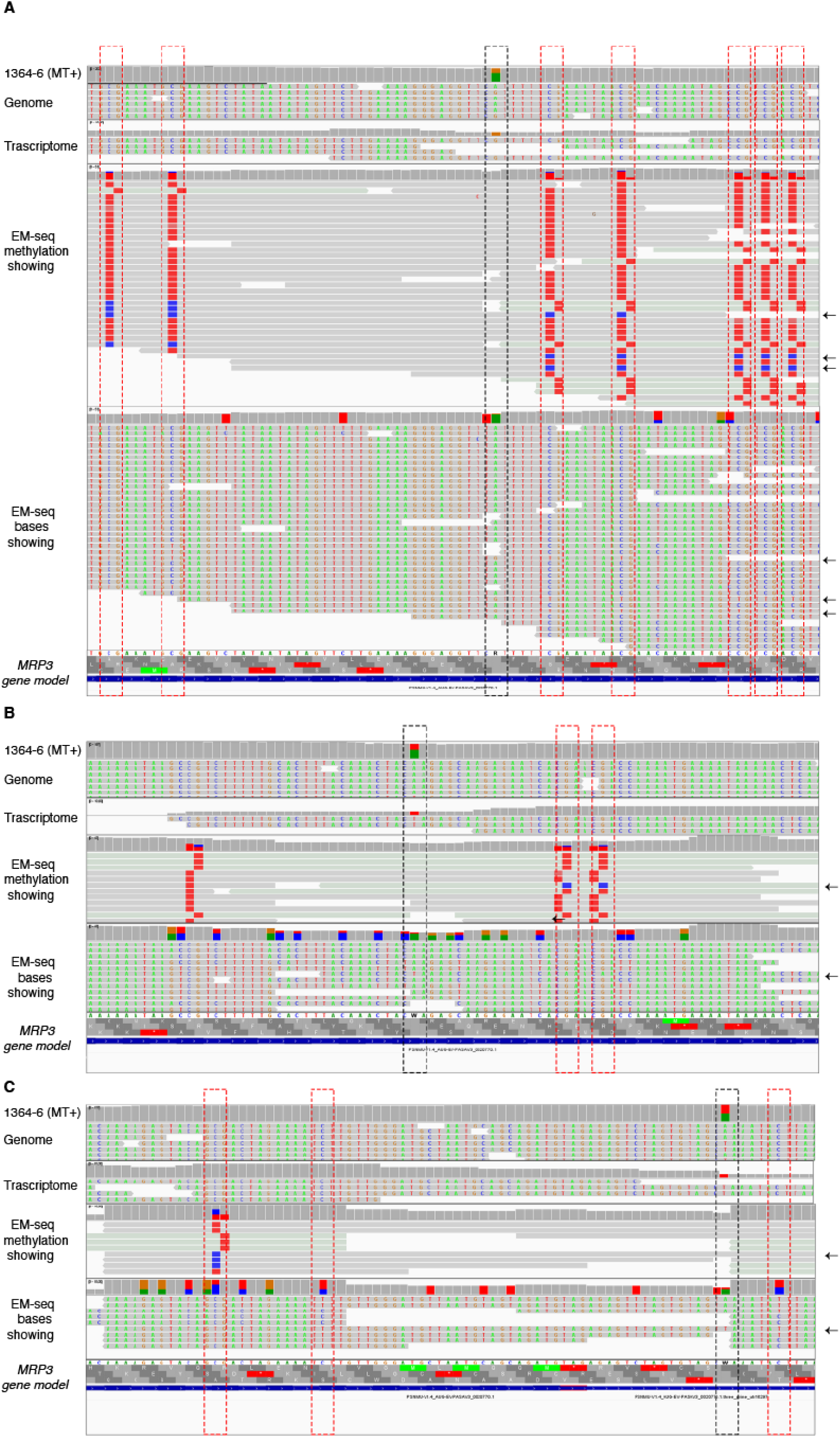
The *MRP3* hypomethylated allele is the transcribed allele. In the three panels, portions of the *MRP3* gene are shown for the MT+ strain 1364-6. In each panel, the top row displays the genome sequence, the second row displays RNA-seq data, the third and fourth rows display the EM-seq data, with the visualization of the methylation profile and of the sequence, respectively, for exactly the same reads. In **A**, a G/A polymorphism is present in the genome (orange and green bars), the transcript is produced from the allele with G, in the EM-seq data the reads with the G, indicated by black arrows on the right, are the ones with hypomethylation. In **B** and **C** similar data are shown for two T/A polymorphisms (red and green bars).

**Figure S4.**
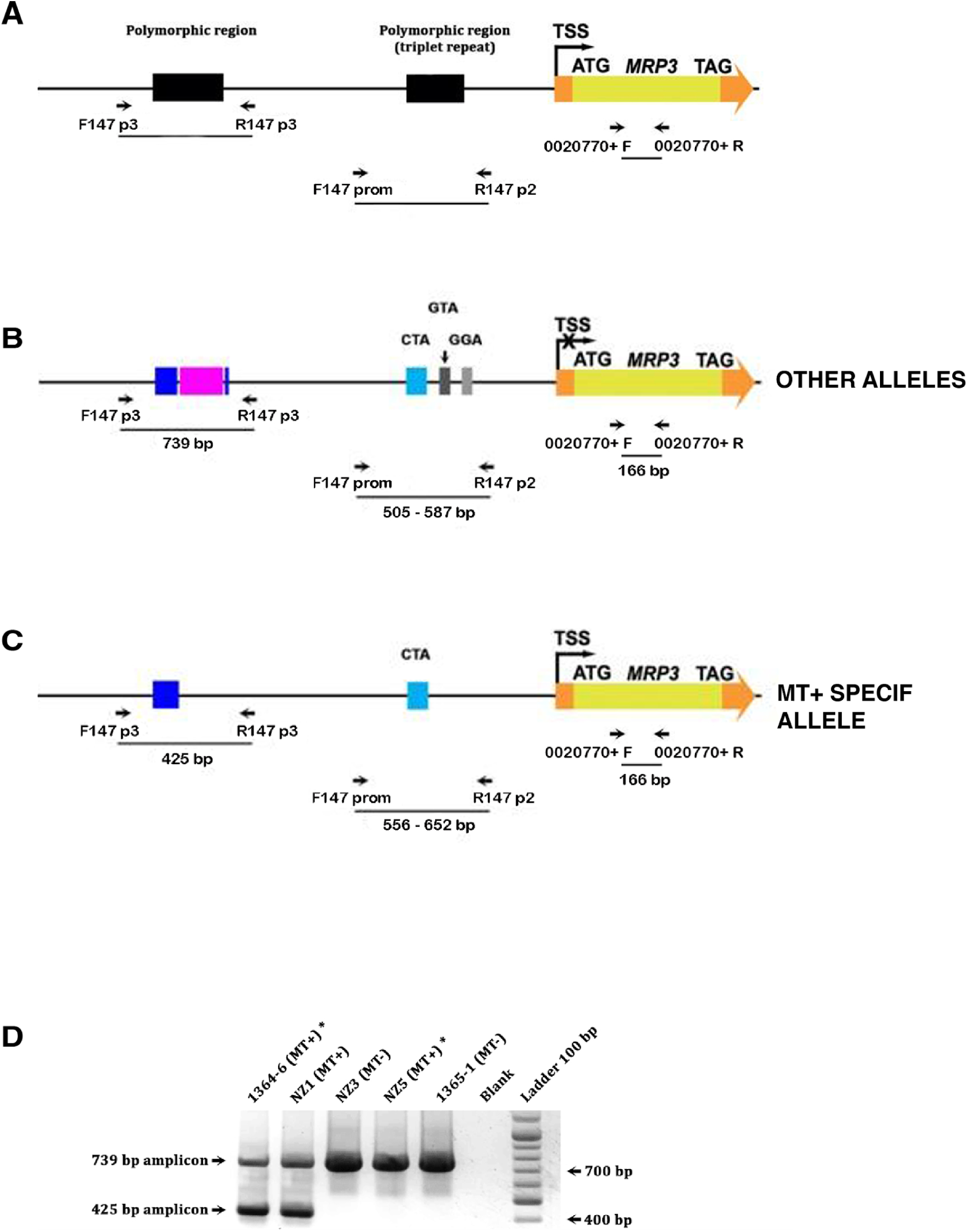
Schematic representation of the*P. multistriata MRP3* alleles. **A,** Schematic representation of the *MRP3* gene region with the primers used to amplify polymorphic regions. **B,** Exemplary alleles identified in different *P. multistriata* populations (*Russo et al., 2018, Russo et al., 2021)*. Different triplets repeats were found in MT+ or MT-strains. In particular, in the alleles of either MT- or MT+ we found from 2 to 33 CTA repeats, from 0 to 16 GTA repeats and from 0 to 16 GGA repeats, while in the MT+ specific allele we found from 26 to 57 CTA repeats and a shorter region upstream (blue box) compared to any other allele (blu and pink boxes). **C,** Electrophoretic gels with amplicons obtained with F147p3/R147p3 primers. Lane 1, 1364-6 (MT+) strain from The Gulf of Naples. Lane 2, NZ1 (MT+). Lane 3, NZ3 (MT-). Lane 4, NZ5 (MT+). Lane 5, 1365-11 (MT)-strain from the Gulf of Naples. Lane 6, blank. Lane 7, GeneRuler 100 bp DNA Ladder (Thermo Scientific™). * denotes strains for which EM-seq data have been produced.

**Figure S5.**
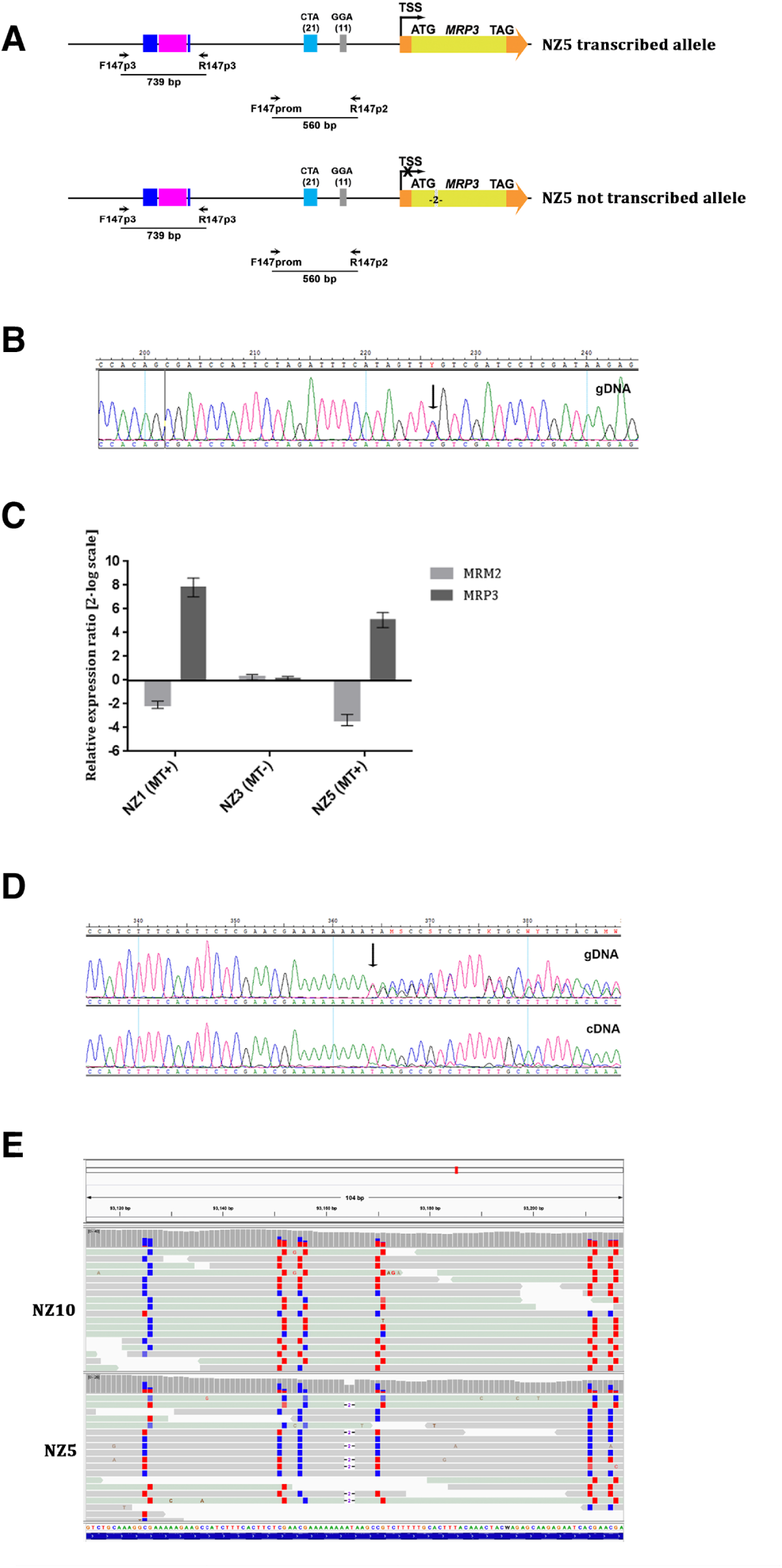
*MRP3* monoallelic expression in NZ5, relative methylation profile and MR expression in NZ strains. **A,** Schematic representation of the NZ5 alleles along with the primers used to sequence the two polymorphic regions. Note that the two alleles differ for a 2 bp indel in the *MRP3* CDS (indicated as -2-). **B,** Electropherograms showing a fragment of the 739 bp amplicon on NZ5 gDNA. A black arrow indicates a SNP that allows to deduce that there are two different alleles. **C**, MT related genes *MRM2* and *MRP3* expression levels in NZ1 (MT+), NZ5 (MT+) and NZ3 (MT-) with respect to the MT-strain 1365-11 used as reference condition and set as zero. **D**, Electropherograms showing the sequence of the *MRP3* CDS fragment amplified from the gDNA (top track) and from the cDNA (bottom track). A black arrow indicates the frameshift due to the 2 bp deletion at the level of the non-transcribed allele. **E,** IGV (Integrative Genome Viewer) EM-seq genome visualization of a *MRP3* CDS fragment on scaffold 147 from NZ10 (MT-) and NZ5 (MT+) strains. By examining the reads, we can observe that in the MT+ NZ5 strain, the non-transcribed allele, with a 2 bp deletion, exhibits methylation indicated by red squares, whereas the transcribed allele, without the 2 bp deletion, is hypomethylated, as indicated by blue squares.

**Figure S6.**
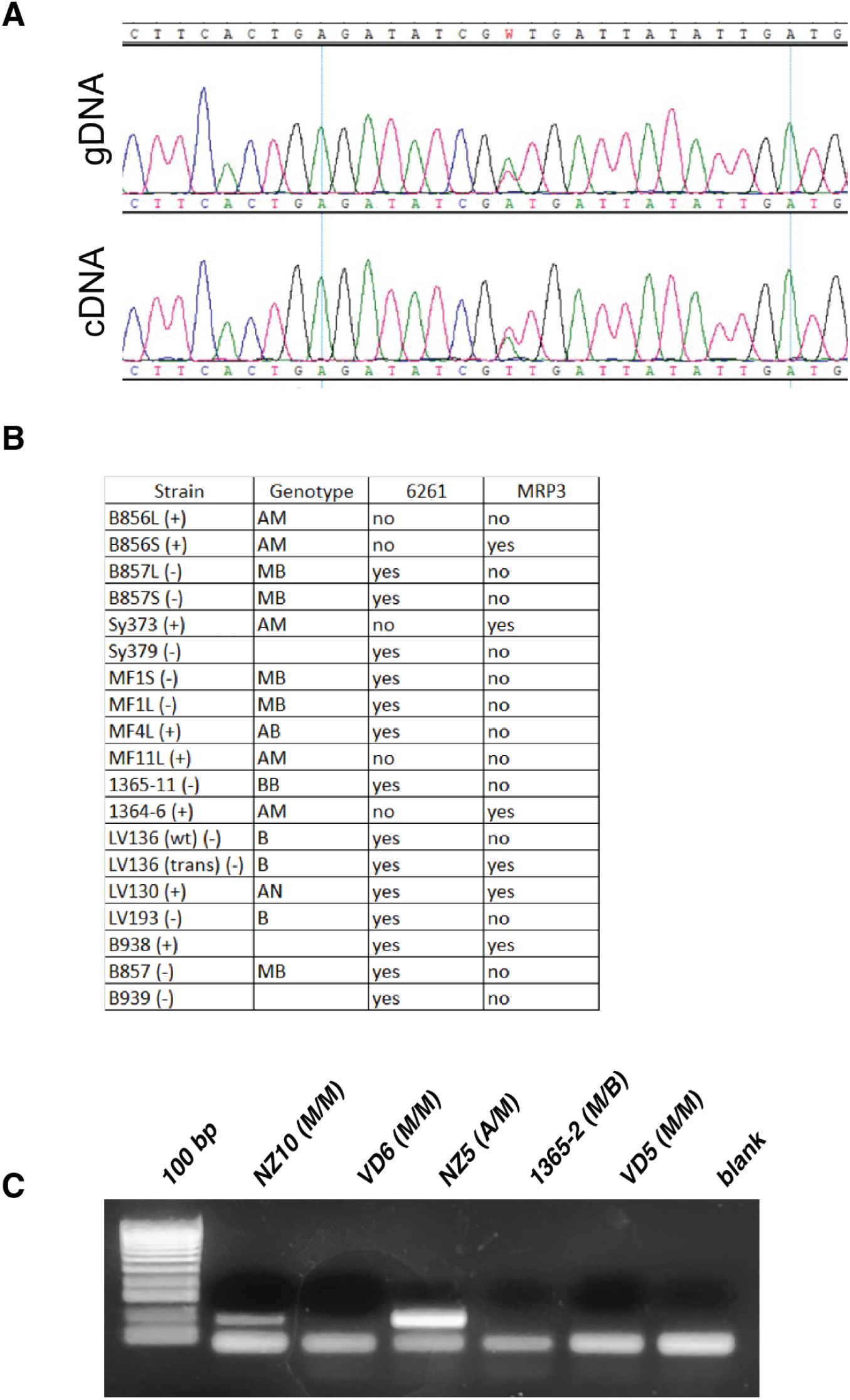
MTase expression. **A,** Electropherograms for the MTase in MF9 gDNA and cDNA show biallelic expression of this gene, as evident from the presence of one SNP. **B,** List of strains in which the MTase and *MRP3* expression was assessed based on available RNAseq datasets (see RNA-seq track at https://bioinfo.szn.it/pmultistriata/). Genotype was assessed by PCR, as described in *Russo et al., 2018 and 2020*. **C,** MTase amplification via RT-PCR on different strains shows that its expression is not correlated to MT nor to allelic variants.

**Figure S7.**
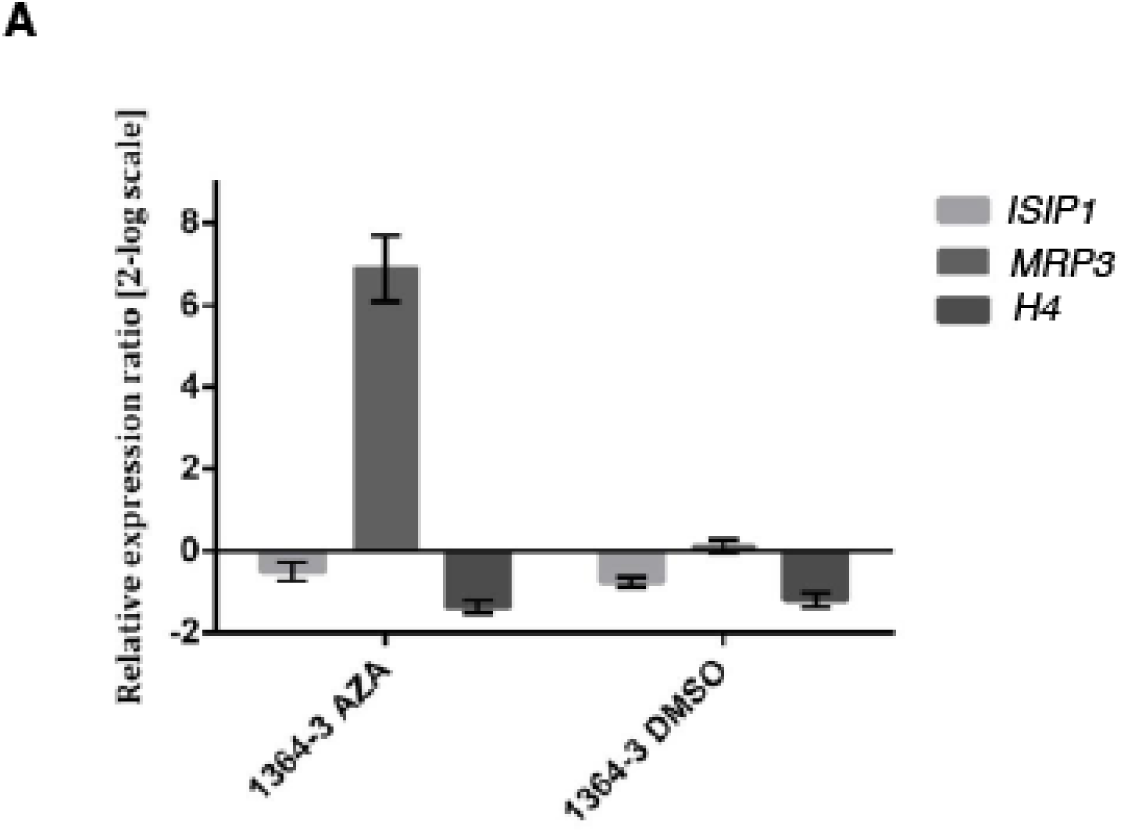
5-AZA effect on transcription of negative control genes. *ISIP1, MRP3* and *H4* transcriptional regulation in the 5-AZA/DMSO treated strains with respect to the untreated 1364-3 MT-strain set as control condition (zero). Data bars represent the mean and standard deviations of data obtained from three different replicates.

**Figure S8.**
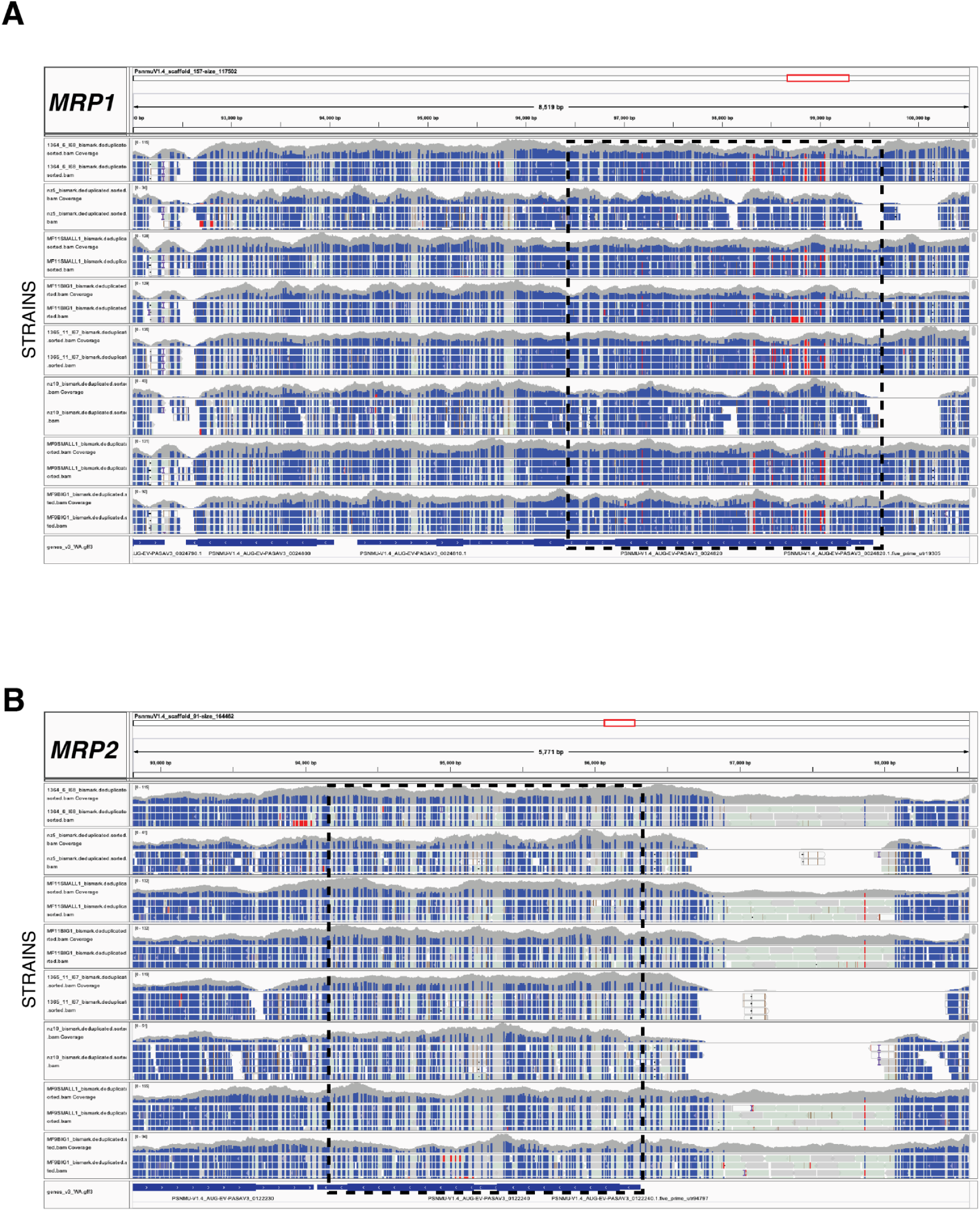
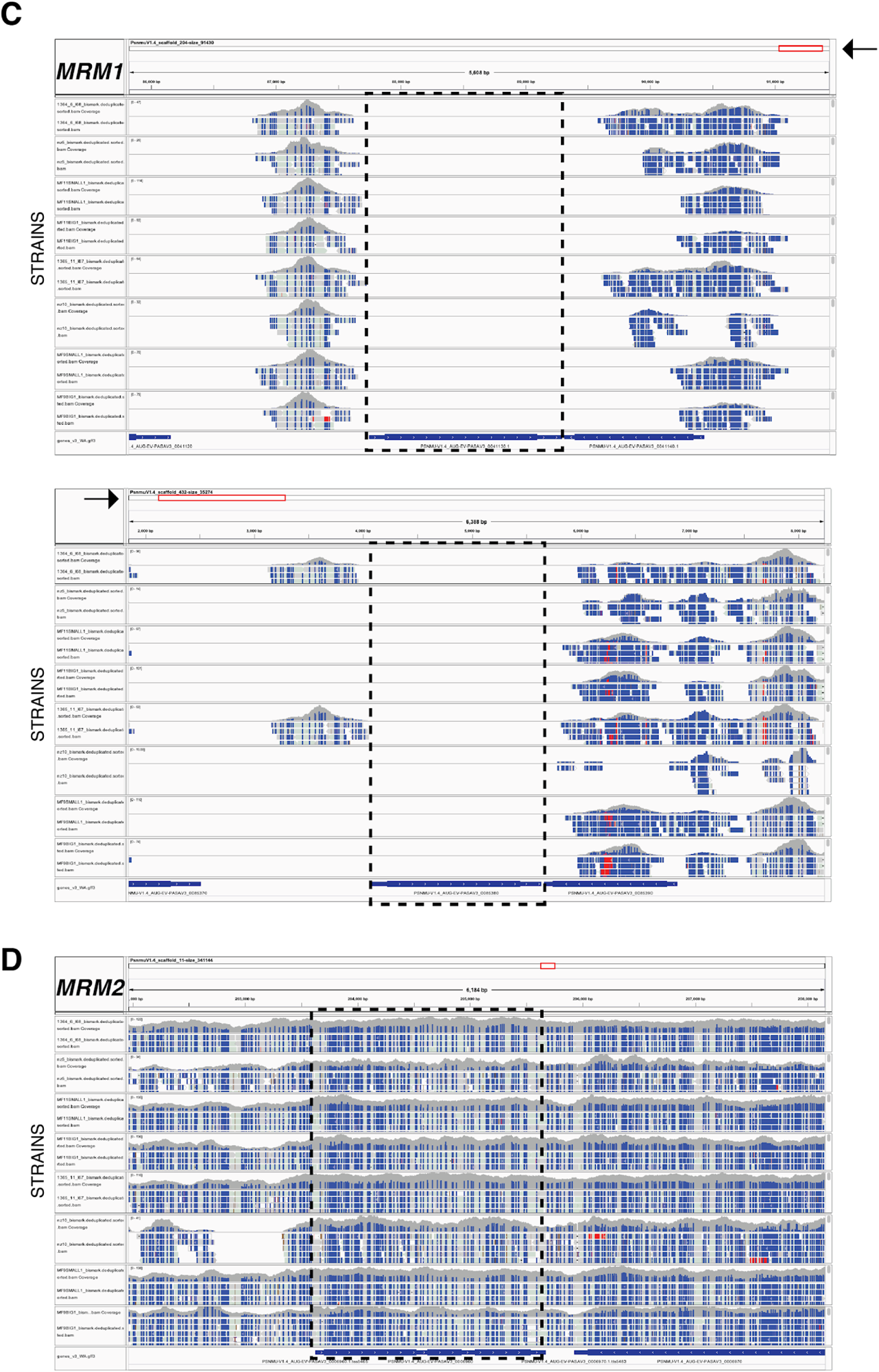
Mating Type Related genes methylation profile.I. IGV visualization of the genomic regions surrounding **A**, *MRP1,* **B***, MRP2,* **C**, *MRM1,* and **D**, *MRM2.* Note that in the current *P. multistriata* reference genome the *MRM1* gene is duplicated at the end of two different, non-overlapping small scaffolds. Black dotted lines highlight the region of interest.

